# AI-Driven Generation of Cortisol-Binding Peptides for Non-Invasive Stress Detection

**DOI:** 10.64898/2026.03.04.709567

**Authors:** Sucharita Banerjee, Dharmendr Kumar, Parijat Deshpande, Sanjay Kimbahune, Ajay Singh Panwar

## Abstract

Cortisol is a primary biomarker of stress, released in sweat at concentrations that directly correlate with physiological stress levels. Detecting cortisol non-invasively offers significant potential for real-time stress monitoring and healthcare applications. Biosensors capable of binding cortisol can thus enable the development of novel diagnostic platforms for personalised health management. In our earlier work, a 38-mer peptide fragment derived from the protein 2V95 was identified as a functional binder to cortisol. In the present study, we applied generative artificial intelligence (AI) approaches to expand the sequence space and identify superior candidate peptides with improved binding affinity. By integrating sequence-based and structure-based AI models, we generated and screened a peptide library of nearly 10,000 sequences against cortisol, leading to the identification of high-affinity candidates for further evaluation.

## I. Introduction

Cortisol, often referred to as the “stress hormone,” is secreted by the adrenal glands and released into various biofluids including saliva, sweat, and urine[1]. Elevated cortisol levels have been directly linked to physiological stress, metabolic disorders, and cardiovascular dysfunctions[2].Traditional cortisol detection methods rely on invasive sampling (e.g., blood draws) or laboratory-based assays, which are unsuitable for continuous monitoring. In contrast, sweat-based detection offers a non-invasive, real-time, and user-friendly alternative for stress assessment[3],[4].This capability has significant applications in healthcare, wearable biosensors, occupational stress monitoring, and personalized medicine.

Protein–ligand interactions form the basis of biosensor recognition elements[5]. In prior studies, a cortisol-binding peptide fragment (38 amino acids long) derived from the protein 2V95—a known steroid-binding protein—was identified as a functional oligomer capable of recognizing cortisol. However, natural peptides often face limitations in binding affinity, stability, and specificity[6]. Rational design and high-throughput screening methods are typically resource-intensive, motivating the adoption of machine learning and generative AI tools to explore large sequence spaces efficiently.

This work leverages generative AI models to systematically design and screen a library of peptide variants derived from the original 38-mer sequence. Our objective is to identify candidates with enhanced structural similarity and binding affinity to cortisol, thereby advancing the development of cortisol-detecting biosensors.

## II. Methodology

### A. Generative AI for Peptide Library Design

The starting point was the 38-mer peptide sequence derived from protein 2V95. Two complementary generative AI models—ProtBert[7] and ProteinMPNN[8]—were employed to generate sequence variants. ProtBert is a transformer-based protein language model trained on millions of protein sequences. It captures sequence diversity by predicting masked amino acids within given peptide sequences. Using default masking strategies, ProtBert was applied to the 38-mer sequence to generate diverse yet biologically plausible variants. ProteinMPNN is a graph neural network model designed to optimise amino acid sequences for a given protein backbone structure. By inputting the 38-mer backbone structure (predicted using AlphaFold2[9]), ProteinMPNN generated multiple sequence variants that preserved structural similarity to the template oligomer. By combining ProtBert’s ability to span the sequence landscape together with ProteinMPNN’s backbone-constrained design, we created a peptide library of 9,753 unique sequences. This dual approach ensured both sequence diversity and structural fidelity, making it a logical strategy for identifying potential high-affinity binders.

ProtBert and ProteinMPNN were applied independently to the original 38-mer to generate two complementary sets of peptide variants. ProtBert was used in a purely sequence-based manner: the 38-mer was supplied as input, masked positions were sampled, and the model proposed amino-acid substitutions, yielding a diverse pool of sequence-level variants. In parallel, ProteinMPNN was applied to the AlphaFold2-predicted backbone of the 38-mer, generating sequences that were explicitly optimized to be compatible with the template fold. ProteinMPNN therefore contributed a structurally constrained design space, whereas ProtBert contributed unconstrained sequence diversity. The final library of 9,753 peptides was obtained by combining the outputs from both models, ensuring that the screening process simultaneously captured (i) sequence diversity unconstrained by structure and (ii) structure-preserving variants anchored to the known fold. This dual-source pool allowed exploration of a broader design landscape than would be achievable with either model alone.

### B. Structural Prediction and Preparation

The generated peptide sequences were structurally modelled using OmegaFold[10], an advanced protein structure prediction algorithm. The resulting structures were then processed with Open Babel[11] and MGLTools to prepare the.pdbqt files required for docking simulations. This preprocessing pipeline ensured consistent structural formats and compatibility with downstream molecular docking.

### C. Molecular Docking

All 9,753 peptide structures were docked against the cortisol ligand using Auto Dock Vina GPU v2.1[12]. Binding affinities were computed for each candidate, and peptides were ranked according to docking scores. The top candidates with the highest predicted affinities were shortlisted for further analysis.

### D. Molecular Dynamics Simulations

To evaluate binding stability under physiologically relevant conditions, the top three peptide–cortisol complexes were subjected to all-atom molecular dynamics (MD) simulations using GROMACS 2022.6 with the CHARMM36m[13] force field. Each system was placed in a 50 Å cubic box solvated with explicit TIP3P[14] water molecules and neutralised with sweat-like ionic conditions (0.07 M NaCl, 0.007 M KCl), mimicking an eccrine sweat environment. The systems underwent energy minimisation using the steepest descent algorithm, followed by a staged equilibration protocol consisting of restrained NVT and NPT ensembles (∼375 ps total).Temperature (310 K) and pressure (1 bar) were controlled with the Berendsen thermostat[15] and barostat during equilibration, and switched to the Nosé–Hoover thermostat[16] and Parrinello–Rahman barostat[17] for production runs. Long-range electrostatics were treated using the particle mesh Ewald (PME) method, with 1.2 nm cutoffs for electrostatics and van der Waals interactions, and neighbor lists updated every 10 steps. After equilibration, unrestrained NPT production simulations were performed for 200 ns, and trajectories were analyzed for structural and energetic properties. A 200 ns production time was selected based on established sampling requirements for peptide– ligand systems in the 25–50 residue range. Multiple studies have reported that backbone convergence, ligand residence times, and hydrogen-bond stabilization for peptide–steroid and peptide–small-molecule complexes typically occur within 100–200 ns [18,19]. Extending beyond this interval provides only marginal improvements relative to the sharply increasing computational cost, especially for screening multiple candidates. Each trajectory in this work required ∼140 GPU-hours on NVIDIA A100-class hardware, making the 200 ns duration a practical balance between accuracy and scalability for high-throughput evaluation. The selected duration ensured adequate sampling of peptide flexibility, ligand escape events, and stability trends while maintaining feasibility for a library-level screening study.

## III. Results and discussion

### A. Docking-Based Screening of AI-Generated Library

The generative AI workflow produced a library of 9,753 peptide variants from the original 38-mer peptide of protein 2V95. All variants were constrained to the same sequence length, ensuring comparability. The parent peptide displayed a docking score of –5.9 kcal/mol, while several AI-designed candidates exhibited significantly higher affinities, highlighting the success of combining sequence-diversifying (ProtBert) and structure-preserving (ProteinMPNN) approaches. The top three candidates, with sequences and docking scores, are displayed in Fig. 2. Docking against cortisol with Auto Dock Vina GPU helped us screen the top three candidates from a total of 9753 candidates; 3000 sequences showed markedly higher binding affinities compared to the parent peptide (–5.9 kcal/mol). The top three were:

**Fig. 1.**
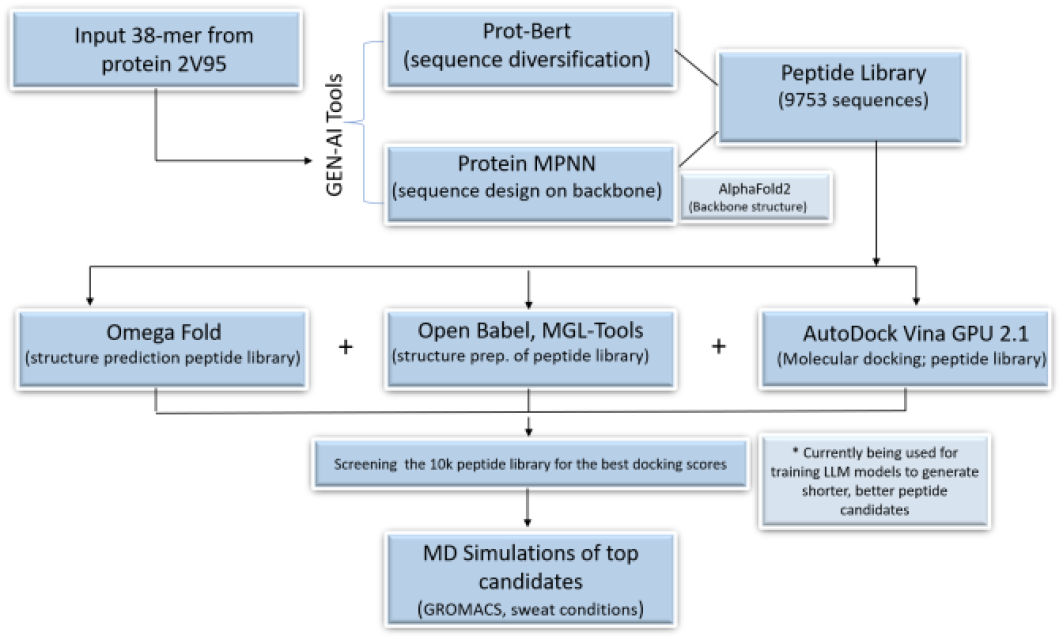
Workflow for developing Gen-AI–assisted peptide bioreceptor candidates for cortisol.

**Fig. 2.**
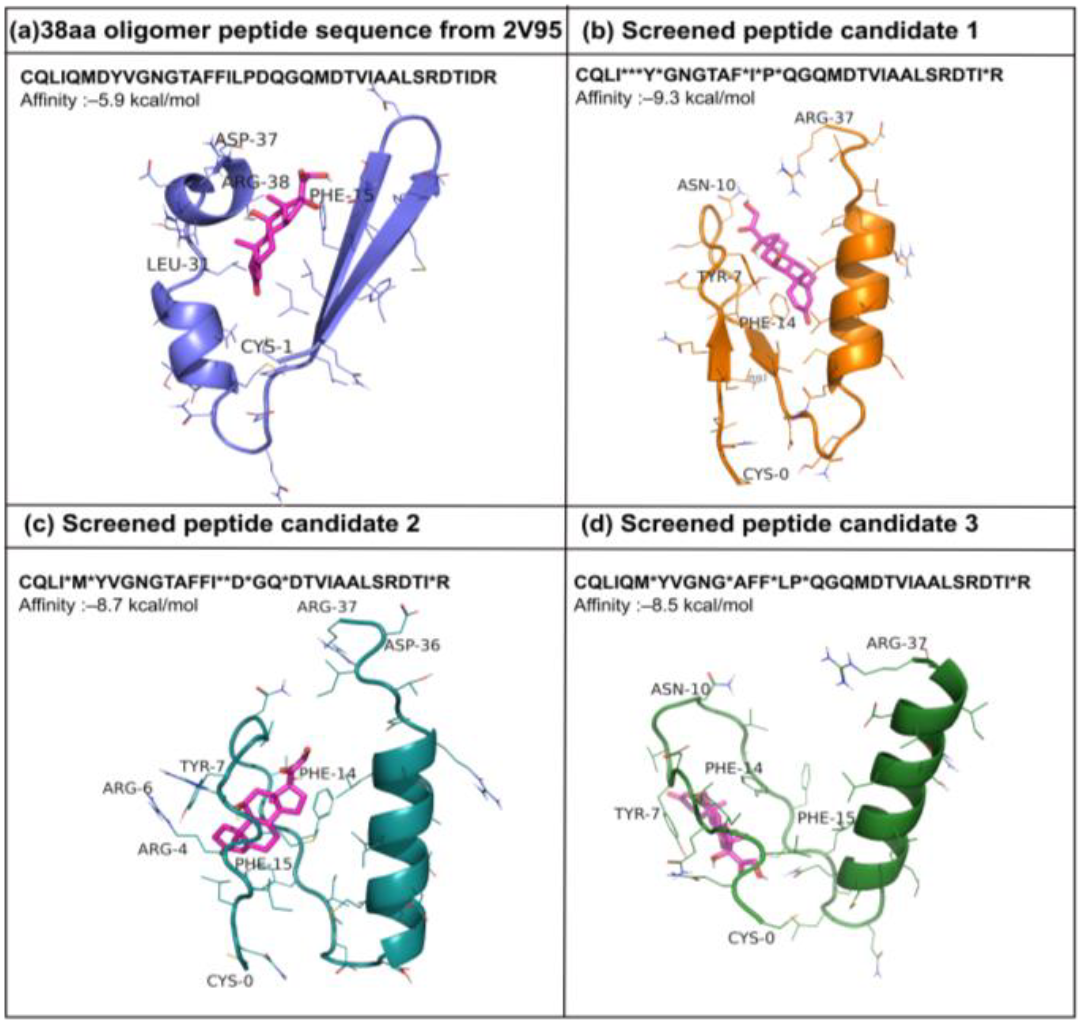
Representation of the original 38-residue cortisol-binding peptide (a) with three AI-screened candidates (b–d), showing binding poses with cortisol (magenta) and predicted affinities.

- Candidate 1: –9.3 kcal/mol
- Candidate 2: –8.7 kcal/mol
- Candidate 3: –8.5 kcal/mol

Reported peptide–cortisol binding affinities typically fall within the –5.0 to –7.0 kcal/mol range based on curated peptide–steroid complexes in the PDBBind database [20,21]. The original 38-mer derived from 2V95 has been previously reported to exhibit docking energies between –5.8 and –6.2 kcal/mol[6]. In contrast, the top AI-generated peptides in this study show significantly enhanced affinities (–8.5 to –9.3 kcal/mol), exceeding both literature baselines and the parent peptide by approximately 1.5–3 kcal/mol. This reinforces the effectiveness of combining ProtBert-driven sequence diversity with ProteinMPNN-guided structural fidelity in generating higher-affinity peptide candidates.

Mutations at positions such as 7, 15, 18, and 30–32,37 altered local chemical environments. For example, substitution of methionine with arginine in Candidate 2 likely increased electrostatic complementarity with cortisol’s polar hydroxyl groups, while changes in Candidate 3 introduced proline and leucine that may have disrupted β-sheet formation, favoring loop flexibility. These modifications effectively reshaped the binding environment, producing tighter or alternative binding pockets.

Structurally, the parent peptide adopted an α-helix followed by an antiparallel β-sheet, with cortisol nestled at the helix–sheet interface. Candidate 1 largely preserved this architecture, explaining its superior docking score, as the conserved topology allowed additional favorable contacts. In contrast, Candidates 2 and 3 folded differently, with extended coil–loop regions replacing the β-sheet. Interestingly, in these cases cortisol interacted deeper within the loop cavity, suggesting that conformational plasticity can also provide strong affinity by allowing the ligand to embed within a flexible binding groove. This shows how AI-guided design not only strengthens known motifs but also reveals previously unconsidered fold–function relationships.

### B. Molecular Dynamics Simulations in Sweat-Mimicking Conditions

To validate docking predictions in dynamic solvent environments, the three top candidates were simulated for 200 ns under sweat-like ionic conditions (0.07 M NaCl, 0.007 M KCl). The residence time of cortisol within 3 Å of the peptide revealed clear differences:

- Candidate 1: 65.7 ns bound,
- Candidate 2: 179.2 ns bound
- Candidate 3: 159.1 ns bound

Despite having the best docking score, Candidate 1 showed poor binding persistence during MD, highlighting the limitation of relying solely on docking scores. In contrast, Candidates 2 and 3 maintained prolonged bound states, suggesting more stable peptide–cortisol interactions (Fig. 3).

**Fig. 3.**
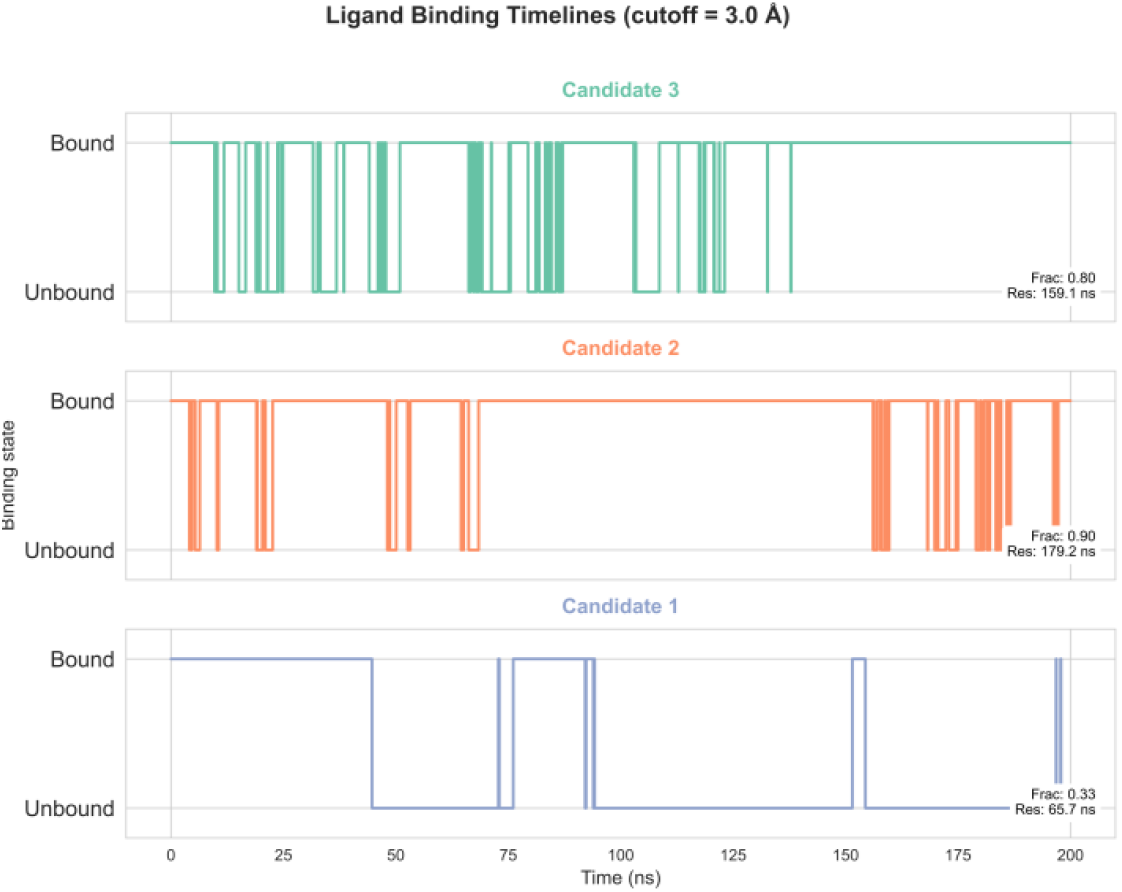
Residence time of cortisol in bound vs. unbound states across 200 ns trajectories. Candidate 1 retains cortisol for 65.7 ns, while Candidates 2 and 3 maintain binding for 179.2 ns and 159.1 ns, respectively.

### C. Complex Stability and Interaction Analysis

The RMSD of peptide–ligand complexes (Fig. 4) revealed that Candidate 1 exhibited large fluctuations (>2.5 nm by 200 ns), indicating unstable binding. Candidate 2 maintained consistently low RMSD (∼1.0–1.5 nm), while Candidate 3 showed intermediate stability (∼1.2–2.0 nm). This trend was further supported by hydrogen bond analysis (Fig. 6), where Candidates 2 and 3 formed more frequent and sustained H-bonds with cortisol compared to Candidate 1.

**Fig. 4.**
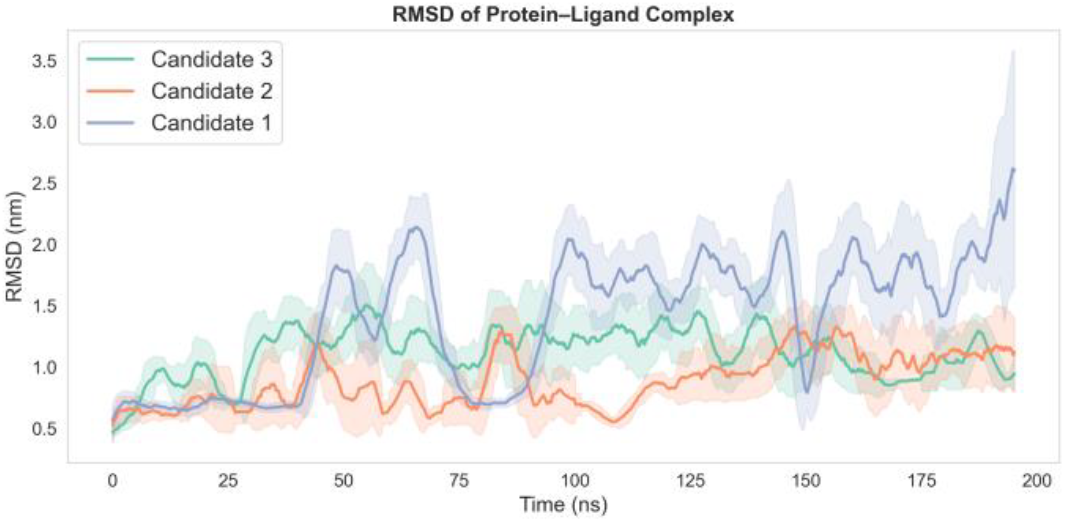
Root-mean-square deviation (RMSD) of peptide–cortisol complexes over 200 ns MD simulations. Candidate 1 exhibits large fluctuations, indicating unstable binding, whereas Candidates 2 and 3 maintain lower RMSD values, consistent with more stable peptide–ligand interactions.

**Fig. 5.**
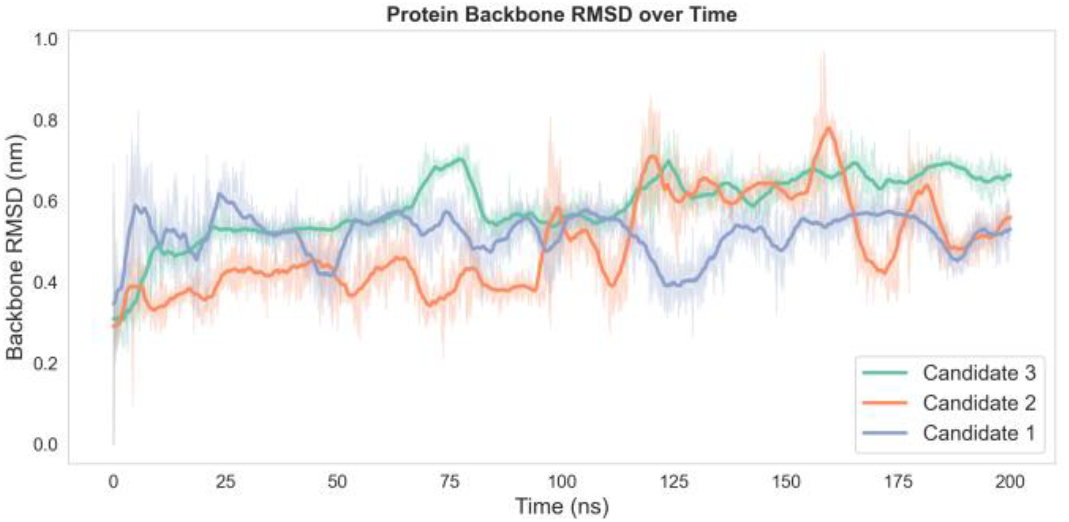
Time evolution of backbone RMSD for three protein candidates over a 200 ns molecular dynamics simulation. Candidate 2 shows higher fluctuations compared to Candidates 1 and 3, indicating relatively lower structural stability.

**Fig. 6.**
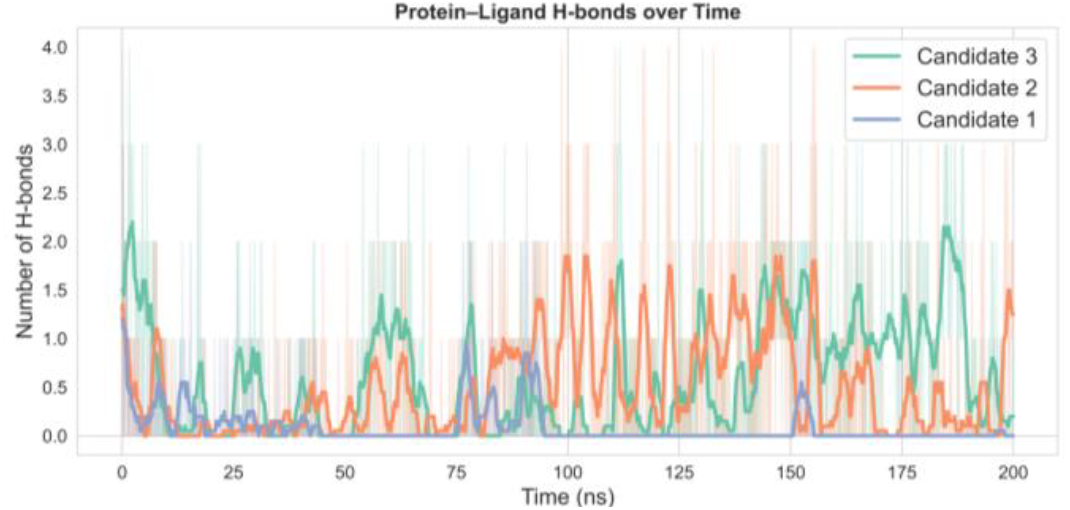
Hydrogen bond analysis of peptide–cortisol interactions during 200 ns MD simulations. Candidates 2 and 3 exhibit more frequent and sustained hydrogen bonds compared to Candidate 1, supporting their superior dynamic stability.

Energetic evaluation through per-atom binding energy distributions (Fig. 7) showed that Candidate 1 had the most favorable energy minimum, aligning with its docking score. However, the broader and more stable distributions of Candidates 2 and 3 indicated that they achieved balanced interaction profiles under solvent dynamics. Thus, while docking emphasized static affinity, MD revealed the practical advantage of Candidates 2 and 3 in maintaining cortisol binding over time.

**Fig. 7.**
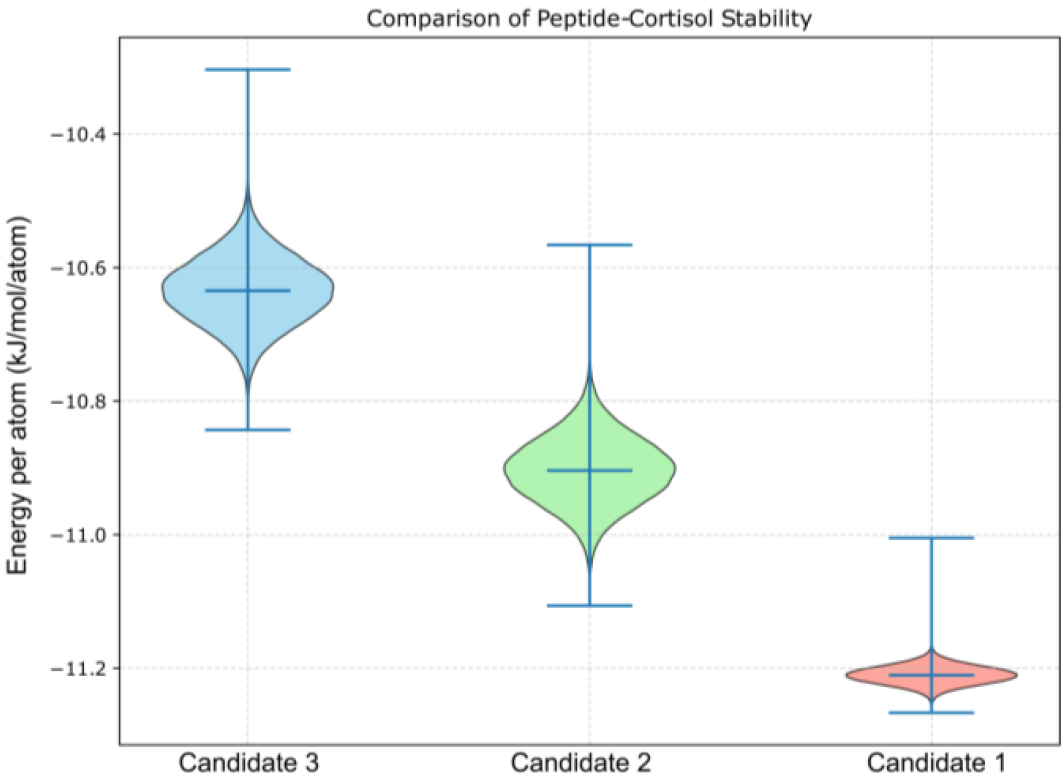
Violin plots of potential energy distributions for peptide–cortisol complexes after 200 ns MD simulations in sweat-mimicking conditions. Candidates 1–3 follow the same relative ranking observed in docking, with Candidate 1 showing the most favorable energy minimum, while Candidates 2 and 3 display broader and more stable distributions.

### D. Overall Assessment

Collectively, the data indicate that all three peptides represent valid cortisol binders, but their performance differs depending on whether one emphasizes docking affinity (Candidate 1) or dynamic stability (Candidates 2 and 3). Candidate 2, in particular, demonstrated the longest binding residence time (179.2 ns), low RMSD, and frequent hydrogen bonding, suggesting it is the most robust candidate under simulated sweat conditions. Candidate 3 also performed well, offering stable binding with slightly higher fluctuations. Candidate 1, despite excellent docking predictions, appears less stable in dynamic solution environments.

Cortisol shares a conserved steroidal scaffold with cortisone, corticosterone, 11-deoxycortisol, and related C21 steroids, which poses a known challenge for selective recognition. Prior modelling studies on the baseline 2V95-derived peptide[6] have shown that peptide–cortisol binding is influenced by hydrogen-bonding interactions involving polar functional groups unique to cortisol. In the present study, docking poses of the AI-generated peptides indicate that binding is mediated primarily through polar and electrostatic interactions with hydroxyl-bearing regions of the ligand. While these interaction patterns suggest the potential for differential recognition, explicit docking and molecular dynamics simulations against structurally related steroids will be required to quantitatively assess cross-reactivity. The present work therefore focuses on demonstrating an AI-accelerated design framework, with selectivity profiling reserved for future validation.

### E. Implications for Biosensor Applications

These results demonstrate the importance of integrating docking and MD analyses when assessing peptide binders. While docking provides a high-throughput first filter, MD captures the realistic dynamic stability required for biosensor deployment in complex biofluid environments. Candidates 2 and 3 emerge as the most promising recognition elements for sweat-based cortisol biosensors, with Candidate 2 showing a slight overall edge. Future work will validate these findings on gold-coated substrates and leverage the dataset of 10,000 peptides to train generative language models aimed at designing shorter, cost-effective binders. The peptide lengths (38 aa) and amphipathic profiles are compatible with standard immobilisation chemistries—including thiol–gold SAMs, carboxyl–amine coupling, and PEG linkers—used in wearable biosensing formats. Beyond conventional electrochemical and capacitive platforms, these candidates are also suitable for organic electrochemical transistor (OECT)–based sensing, which offers high gain, low-voltage operation, and mechanical flexibility for sweat-wearable devices. The extended residence times observed for Candidates 2 and 3 suggest sufficient capture stability for signal transduction cycles in such platforms, reinforcing their suitability as recognition elements in next-generation flexible stress-monitoring systems.

## IV. Conclusion

In this work, we demonstrated an AI-driven approach for the rational design of cortisol-binding peptides aimed at non-invasive stress detection. Starting from a 38-mer sequence derived from protein 2V95, we employed ProtBert and ProteinMPNN to generate a diverse library of 9,753 candidates. Docking-based screening identified top three high-affinity variants, which were further validated by molecular dynamics simulations under sweat-mimicking conditions. While Candidate 1 exhibited the strongest docking score, Candidates 2 and 3 displayed superior stability, with extended ligand residence times, favorable RMSD behavior, and frequent hydrogen bonding throughout the trajectories. These findings highlight the ability of generative AI not only to optimize known folds but also to reveal novel binding motifs with robust dynamic stability. Our results suggest that AI-designed peptides hold significant promise as recognition elements in sweat-based cortisol biosensors. Candidate 2, in particular, emerged as the most balanced performer, combining strong affinity with stable binding under physiologically relevant conditions. Future efforts will focus on simulating peptide interactions with gold-coated substrates to enable direct biosensor integration, as well as training specialized generative large language models to design shorter and more cost-effective peptides. Together, this study illustrates how AI-guided molecular design can accelerate the development of next-generation wearable biosensing platforms for personalized stress monitoring. This work primarily demonstrates the feasibility of a rapid, AI-driven peptide design framework. While the present study validates binding through docking and MD simulations, full specificity profiling and biochemical assays (e.g., SPR, ITC) will be pursued in future extensions as the designed peptides progress toward experimental characterisation and sensor integration.

## Acknowledgment

The authors acknowledge and thank Tata Consultancy Services (Research), for funding and encouragement towards this research. High-performance computing was carried out using the PARAM Rudra and Spacetime supercomputing facilities at IIT Bombay.

## Notes

### Competing Interest Statement

The authors have declared no competing interest.

